# *Dickeya colocasiae* sp. nov. isolated from wetland taro, *Colocasia esculentum*

**DOI:** 10.1101/2022.01.14.476417

**Authors:** Gamze Boluk, Shefali Dobhal, Dario Arizala, Anne M. Alvarez, Mohammad Arif

## Abstract

Bacterial pathogens identified as *Dickeya* sp. have recently been associated with a corm rot of wetland taro on Oahu, Hawaii, but the species designation of these strains was unclear. A Gram-negative, pectinolytic bacterial strain PL65^T^ isolated from an infected taro corm was subjected to polyphasic analysis to determine its genomic and phenotypic characteristics. Multi-locus sequence analyses (MLSA) based on five housekeeping genes (*dnaA, gapA, gyrB, atpD*, and *purA*) revealed that *Dickeya zeae* and *D. oryzae*, were the closest relatives. Phylogenetic analysis based on 463 core gene sequences clearly showed two potentially new species within *Dickeya oryzae. In silico* DNA– DNA hybridization value of strain PL65^T^ with 12 Type strains of *Dickeya* species was <68%. Average nucleotide identity (ANI) analysis revealed that PL65^T^ was at the margin of the species delineation cut-off values with a 96% ANI value. The metabolic profile of strain PL65^T^ using BIOLOG differentiated it from the type strains of all other known species of *Dickeya*. Based on the results of genome-to-genome comparisons and phenotypic data presented in this report, we propose establishment of a new species, *Dickeya colocasiae* sp. nov. with strain PL65^T^ as the type strain (ICMP 24361^T^).

## Introduction

Bacteria in the genus *Dickeya* are rod-shaped Gram-negative bacteria belonging to the family *Pectobacteriaceae* in the order *Enterobacterales* (Adeolu et al., 2016). The genus *Dickeya* produces plant cell wall-degrading enzymes (CWDEs) that cause maceration of plant tissues in field and during post-harvest storage (Collmer and Keen, 1986). *Dickeya* species also have been isolated from irrigation water (Parkinson et al., 2014; Sueno et al., 2014; Hugouvieux-Cotte-Pattat et al., 2019; Oulghazi et al., 2019). Members of the genus *Dickeya* have a broad host range, infecting both monocot and dicot plants, including vegetable crops and ornamental plants (Ma et al., 2007; Charkowski et al., 2012). Mansfield et al. reported *Dickeya* among the top ten most important bacterial plant pathogens based on their economic and scientific impact (Mansfield et al., 2012).

The genus *Dickeya* was first described in 2005 by reclassifying *Pectobacterium chrysanthemi* (Brenner et al. 1973), (formerly *Erwinia chrysanthemi* (Burkholder et al. 1953)) into six species, namely *D. chrysanthemi, D. paradisiaca, D. zeae, D. dianthicola, D. dadantii*, and *D. dieffenbachiae* (Samson et al., 2005). In 2012, *D. dieffenbachiae* was reclassified as subspecies of *D. dadantii* based on DNA-DNA hybridization and multi-locus sequence analysis (Brady et al., 2012). Later, additional species were discovered; *Dickeya solani* was described as the causal agent of severe disease outbreaks of potatoes in Europe (van der Wolf et al., 2014) and *Dickeya fangzhongdai* was described as a novel species, initially isolated from pear trees in China (Tian et al., 2016). Three new species were identified in water samples: *D. aquatica* isolated from freshwaters in England and Finland (Parkinson et al., 2014), *D. lacustris* from lakes in France (Hugouvieux-Cotte-Pattat et al., 2019), and *D. undicola* from water samples collected in Malaysia and France (Oulghazi et al., 2019). More recently, two additional *Dickeya* species have been described: *D. oryzae* isolated from roots of infected rice (Wang et al., 2020) and *D. poaceiphila* isolated from sugarcane (Hugouvieux-Cotte-Pattat et al., 2020). Currently, this genus encompasses 12 recognized species. The main events in the taxonomic history of *Dickeya* are summarized in the timeline (**Figure 1**).

**Figure 1.**
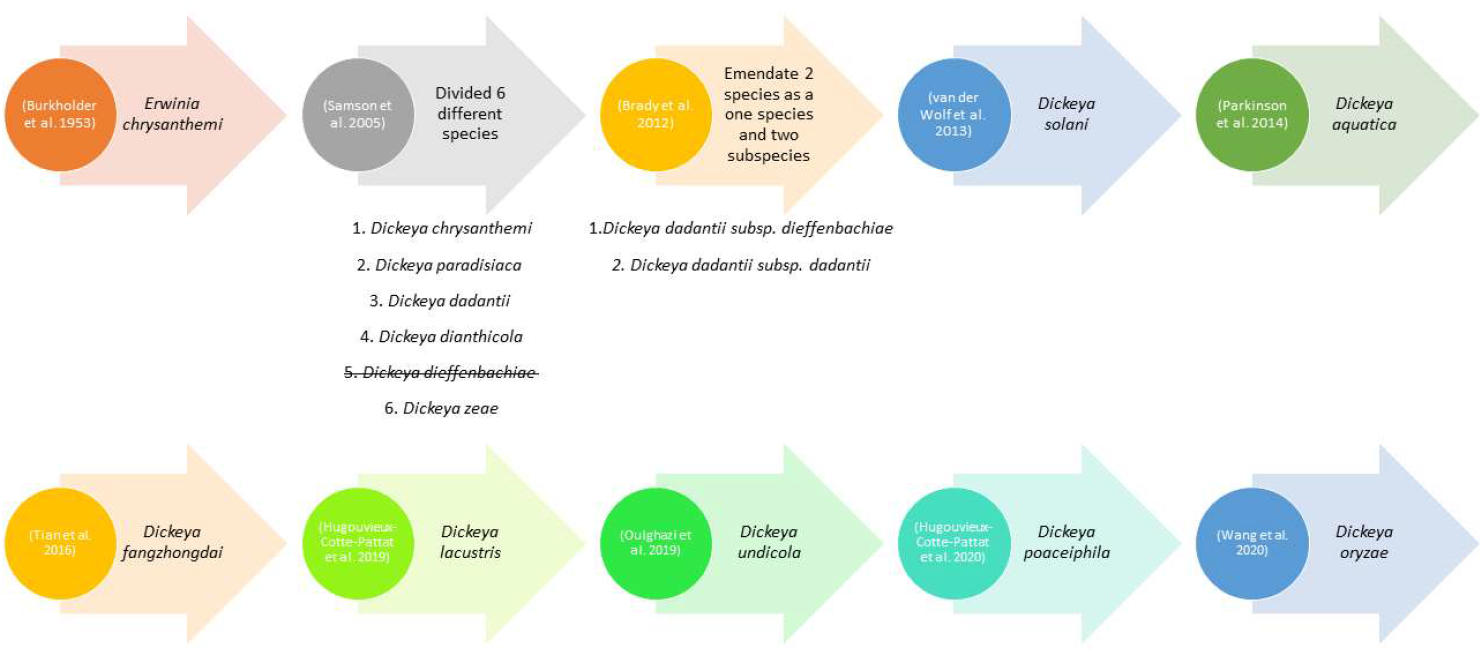
Timeline highlighting the main events based on taxonomic description of soft rot pathogen within the genus *Dickeya*.

In the current study five strains, PL65^T^ isolated from an infected taro corm (Boluk et al., 2021); A5428, A5511, A5432, and A5612, isolated from irrigation water on nearby farms on Oahu, Hawaii (Boluk et al., 2020). These strains belong to the genus *Dickeya* but differed from previously characterized species with respect to several important phenotypic and genomic characteristics (**Table 1**). We propose establishment of a novel species with strain PL65^T^ as the type strain for *Dickeya colocasiae*. We further propose that strain CE1 isolated in 2017 from Canna lily in China (**Table 1**) (Yang et al., 2019) now be reclassified as *Dickeya colocasiae*, based on its genotypic characteristics.

**Table 1.** Detail information about the six strains of novel species of *Dickeya colocasiae*

| Strain | Collected Origin, Location and Year | Isolated by |
| --- | --- | --- |
| PL65<sup>T</sup> | *Colocasia esculenta*, Hawaii, USA, 2018 | Gamze Boluk and Mohammad Arif |
| CE1 | *Canna edulis*, Guangdong, China, 2017 | Jingxin Zhang |
| A5428 | Irrigation water, Hawaii, USA, 2007 | Anne M Alvarez |
| A5432 | Irrigation water, Hawaii, USA, 2007 | Anne M Alvarez |
| A5511 | Irrigation water, Hawaii, USA, 2006 | Anne M Alvarez |
| A5612 | Irrigation water, Hawaii, USA, 2008 | Anne M Alvarez |

## Isolation

Strain PL65^T^ was isolated in 2018 from a wetland taro corm during harvest near in Honolulu, Hawaii (Boluk et al., 2021). Small (1 mm^2^) sections of infected tissue were sterilized for 1 min in 0.6% sodium hypochlorite, rinsed three times with sterile water, triturated in a sterile 1.5 ml centrifuge tube containing 100 μl sterile water, and streaked onto crystal violet pectate (CVP) agar plates (Boluk et al., 2021). Pectolytic colonies that formed pits on CVP, were re-streaked onto dextrose peptone agar plates to obtain single colonies (Boluk et al., 2021). Strains A5428, A5511, A5432, and A5612 were isolated in 2006-2008 from irrigation water in and around pineapple fields showing high incidence of pineapple heart rot in Hawaii between (Sueno et al., 2014). Ten ml samples of irrigation water were passed through 0.45-μm Millipore filters; the filters were placed face up onto the surface of petri plates containing MS medium (Miller-Schroth, 1972) and incubated at 28 °C (Sueno et al., 2014). Bacterial colonies were re-streaked onto a peptone-glucose medium containing tetrazolium chloride plates to differentiate colonies (Sueno et al., 2014). Single colonies were selected, restreaked, and preserved at −80°C in peptone broth containing (30% glycerol, v/v) (Boluk et al., 2021).

## Phylogenetic Analyses

The taxonomic position of strain PL65^T^ was refined using a multi-locus sequence analysis (MLSA) (Glaeser and Kämpfer, 2015). The MLSA based on five housekeeping genes (dnaA, *gapA, gyrB, atpD*, and *purA*) was used for phylogenetic analysis (Boluk et al., 2020). The concatenated nucleotide sequences of *dnaA, gyrB, gapA, atpD*, and *purA* genes of 39 strains were retrieved from the NCBI GenBank genome database (**Table S1**) and aligned using MEGAX software. Strains belonging to the species of *Pectobacterium* were used as outgroups. A phylogenetic tree was reconstructed using the Neighbor-joining algorithm, and the stability of the topology of the phylogenetic tree was evaluated by performing bootstrap analysis based on 1000 replications (Felsenstein, 1985; Saitou and Nei, 1987). Strain PL65^T^, A5428, A5511, A5432, A5612, and CE1 possessed very similar sequences with a high bootstrap value and formed a separate phylogenetic lineage from *D. zeae, D. oryzae*, and Ech586 based on five housekeeping genes. Strain Ech586 was also separated from the other species (**Figure 2**). It was suggested that PL65^T^ and Ech586 should be reclassified as two novel species within the genus *Dickeya*.

**Figure 2.**
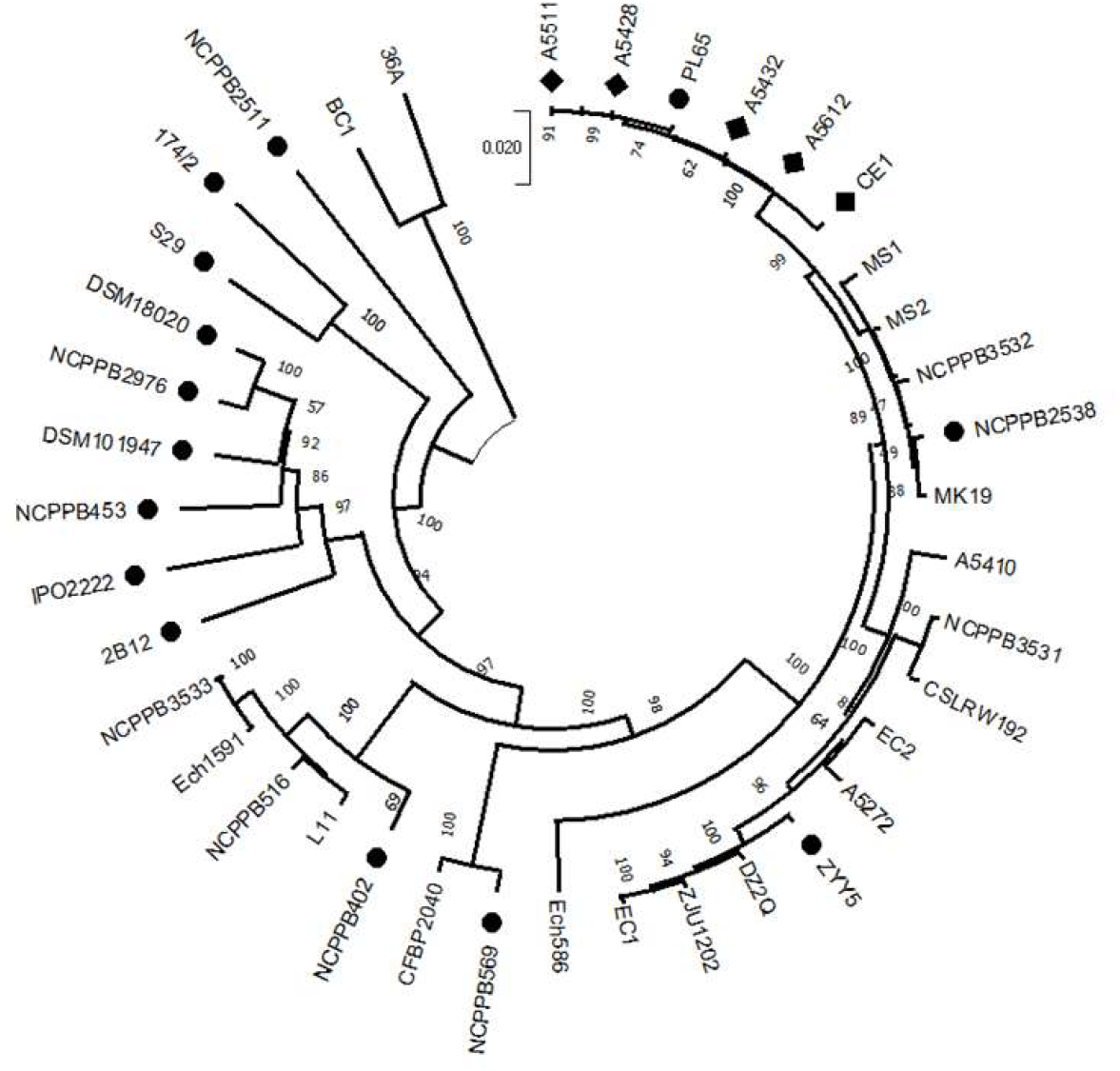
Concatenated phylogenetic analysis using 5 housekeeping genes: chromosomal replication initiator protein DnaA (*dnaA*), DNA gyrase subunit B (gyrB), Glyceraldehyde-3-phosphate dehydrogenase A (gapA), ATP synthase subunit beta (*atpD*), and Adenylosuccinate synthetase (*purA*). Numbers on the nodes represent bootstrap values and presented as a percentage of 1,000 replicates. Neighbor-Joining (NJ) phylogenetic tree of *Dickeya*. *Pectobacterium* species as an out-off group.

## Genome Features

The genome of the novel strain PL65^T^ (CP040817) has been reported and analyzed in our previous comparative genomic analysis of novel *D. zeae* strains isolated from pineapple and taro (Boluk et al., 2021). The results of a comparative genomics study indicated that strain PL65^T^ (CP040817) should be classified as a novel species. The genome of PL65^T^ (CP040817; one complete genome, no plasmid, 701x depth of genome coverage) is 4,749,968 bp and has a DNA G+C content of 53.6% (Boluk et al., 2021). The genome consists of 4,095 protein-coding genes organized within a single replicon (Boluk et al., 2021).

## Comparative Genomic Analysis

The delineation of *D. colocasiae* as a novel species was supported by calculating the digital DNA-DNA hybridization (dDDH) and the average nucleotide identity. The dDDH value of 70% corresponded to ANI values of 95–96% and conserved DNA values of 69% are being used as a boundary for species delineation (Goris et al., 2007). The dDDH or genome-to-genome distance values among 18 genomes were calculated *in silico* using the webserver Genome-to-Genome Distance Calculator (GGDC) version 2.1 with the formula 2 and the BLAST+ alignment criteria (Meier-Kolthoff et al., 2013). The dDDH value between *D. zeae* and *D. colocasiae* was 68.5%, below the recommended cut-off for species delineation 70% (**Figure 3**). Additionally, The Average Nucleotide Identity (ANI) based on the Nucleotide MUMmer algorithm (ANIm) was calculated using the JSpecies Web Server with default parameters (Richter and Rosselló-Móra, 2009). These analyses were performed using the genomes of the type strains of *Dickeya* species. The ANIm values between *D. zeae* and *D. colocasiae*. Ranged between 96.15–96.32 %, which is above the general cut-off for species delineation (95–96 %) (Goris et al., 2007; Richter and Rosselló-Móra, 2009; Kim et al., 2014; Chun et al., 2018; Pasanen et al., 2020; Wang et al., 2020) (**Figure 3**). However, these ANIm values were based on a low conserved DNA percentage, 85.83-88.75%, between the two species (Pasanen et al., 2020) (**Table 2**, **Figure 4**, and **Figure S1**). Moreover, the *D. colocasiae* genomes were more than 98% in ANIm and showed an alignment percentage of 93.12%. The analysis of ANIm, conserved DNA percentage and dDDH showed that *Dickeya colocasiae* genomes resemble *D. zeae* but differ in several important aspects from the other genomes.

**Figure 3.**
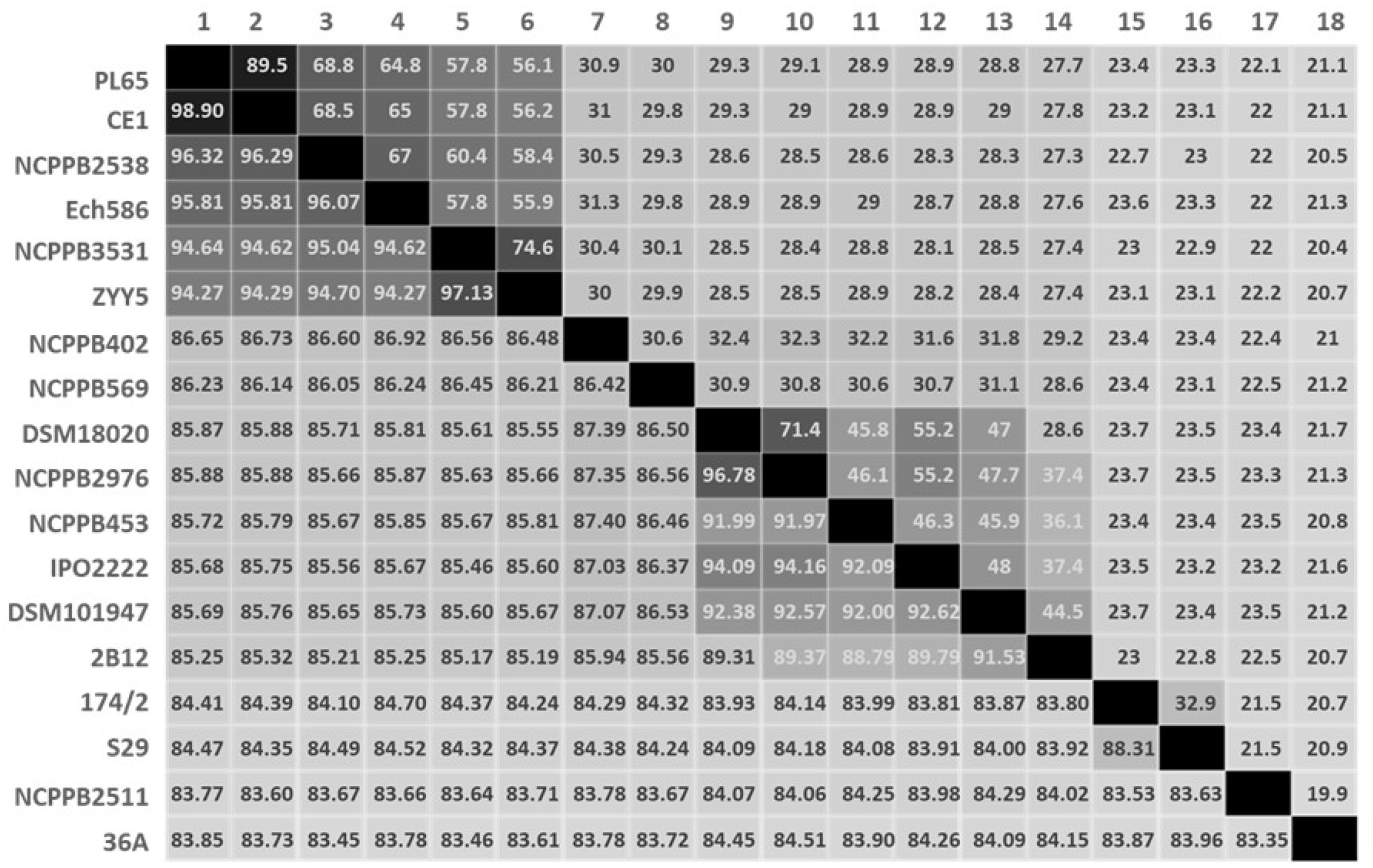
Pairwise heatmap based on the average nucleotide identity (ANI) and digital DNA-DNA hybridization (dDDH). Both ANI and dDDH values are represented in terms of percentage between the reported fifteen *Dickeya* species and one *Pectobacterium* species. Cut-off values for species delineation are 95~96 and 70% for ANI and dDDH, respectively. 1. PL65; *Dickeya colocasiae*, 2. CE1; *D. colocasiae*, 4. Ech586; *D. zeae*, 3. NCPPB 2538; *D. zeae*, 5. NCPPB 3531; *D. oryzae*, 6. ZYY5; *D. oryzae*, 7. NCPPB 402; *D. chrysanthemi*, 8. NCPPB 569; *D. poaceiphila*, 9. DSM 18020; *D. dadantii*, 10. NCPPB 2976; *D. dadantii*, 11. NCPPB 453; *D. dianthicola*, 12. IPO 2222; *D. solani*, 13. DSM 101947; *D. fangzhongdai*, 15. 174/2; *D. aquatica*, 14. 2B12; *D. undicola*, 16. S29; *D. lacustris*, 17. NCPPB 2511; *D. paradisiaca* and 18. 36A; *Pectobacterium atrosepticum*.

**Figure 4.**
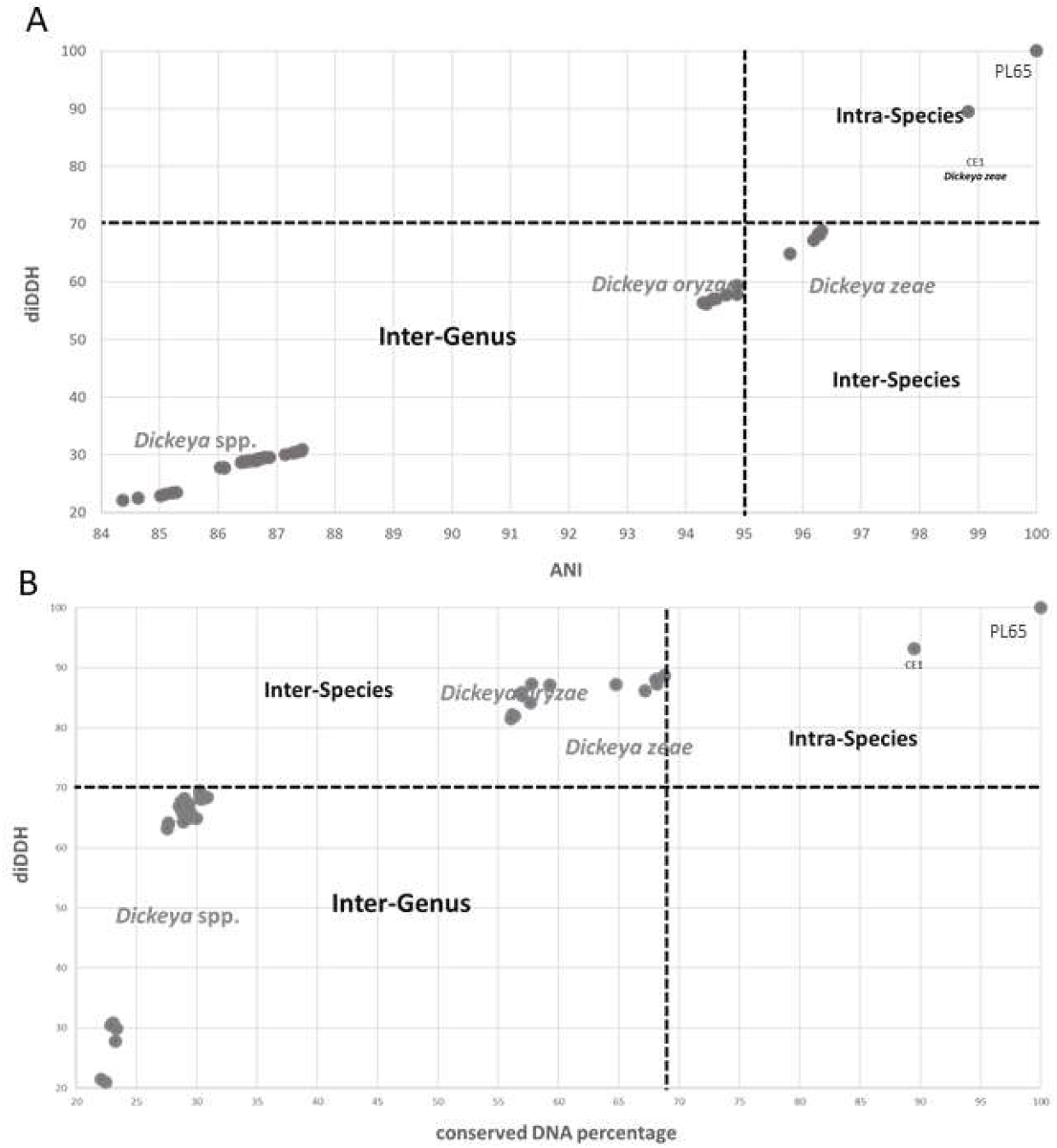
Relationship between digital DNA-DNA hybridization (dDDH) values and average nucleotide identity (ANI) and conservation. Each filled circle shows the value for dDDH between two strains (y-axis), (A) plotted against the ANI of the conserved genes between the strains, and (B) the percentage of conserved DNA between the strains.

**Table 2.** Genomic comparisons of *Dickeya colocasiae* strains with *Dickeya zeae, D. oryzea* and type strains of known *Dickeya* species. The number of strains used in each comparison are shown in the parentheses.

|  | GGDC | ANI | AP |
| --- | --- | --- | --- |
| SPECIES | PL65 and CE1 | PL65 and CE1 | PL65 and CE1 |
| PL65 and CE1 | 89.5 | 98.90 | 93.12 |
| Ech586 | 65-64.8 | 95.81-95.81 | 87.13-86.92 |
| *Dickeya zea* (5) | 68.5-68.5 | 96.15-96.32 | 85.83-88.75 |
| *Dickeya oryzae* (9) | 56.1-57.8 | 94.27-94.86 | 81.21-87.14 |
| *Dickeya* species (11) | 31.3-21.5 | 86.48-83.53 | 21.48-68.51 |

Genomic data used to perform the phylogenomic analysis were based on 463 core gene sequences to enhance the resolution and confidence. Strains PL65^T^, CE1, Ech586, and the other related *D. zeae*, *Dickeya* species, and *P. atrosepticum* were included for core gene analyses (**Table S2**). The 39 genomes were retrieved from NCBI and reannotated with the Prokka software tool (Seemann, 2014). The pan-genomes were analyzed in the Roary pipeline, and the core genome alignment was generated using multi-FASTA alignment based on 463 core genes generated using MAFFT (Page et al., 2015). Later, maximum likelihood trees based on the previously identified core genes were constructed using RAxML-NG (Next Generation) (Kozlov et al., 2019). The maximum likelihood trees were produced for viewing and editing using FigTree v1.4.4 (Rambaut, 2010). The topology of the phylogenetic tree displayed the separation of the *D. zeae* strains, and, in this case, PL65^T^ and CE1 constituted a well-supported cluster (**Figure 5**).

**Figure 5.**
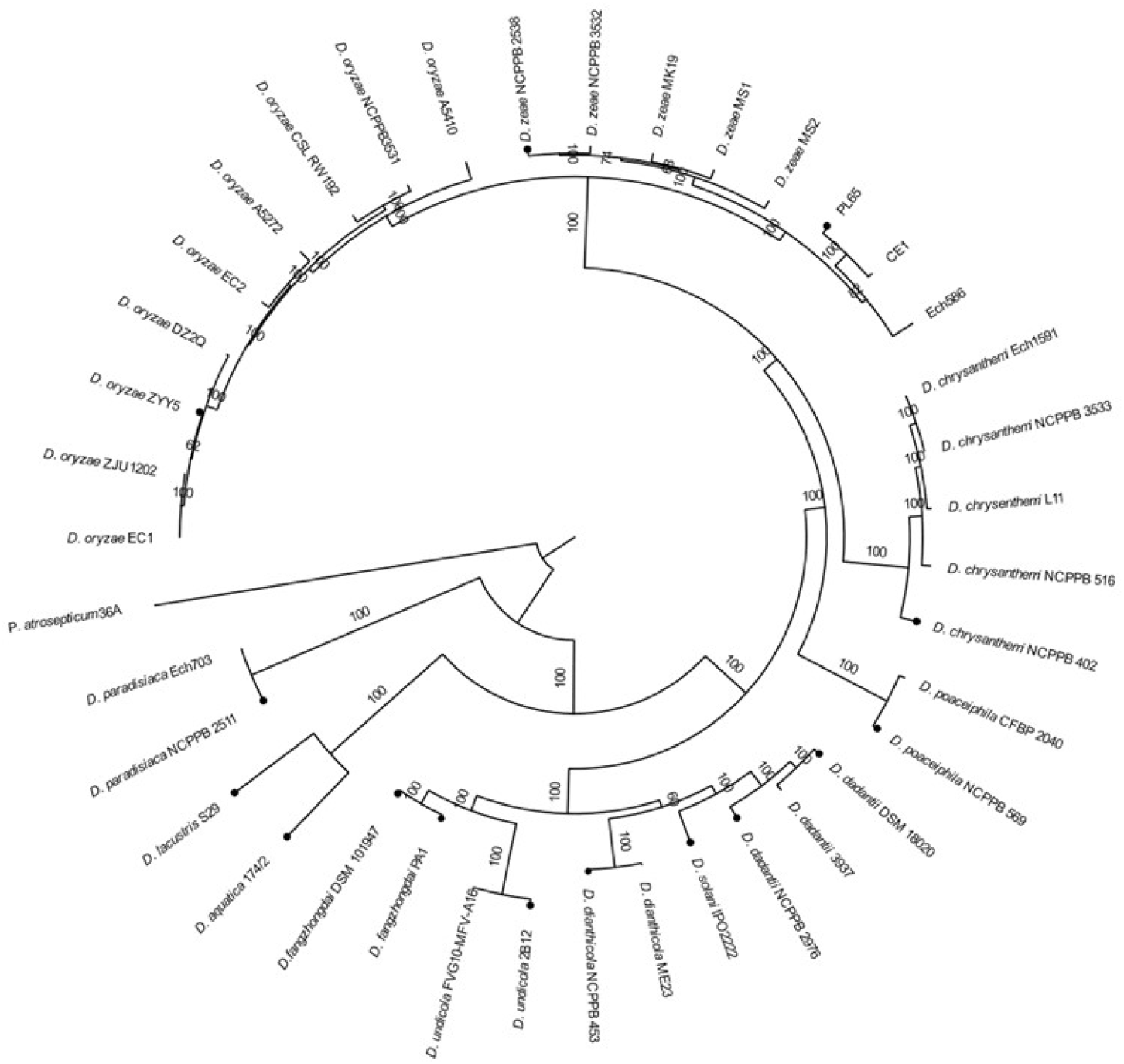
Maximum-likelihood phylogenetic analysis of *Dickeya* species. *Pectobacterium* species was used as an outgroup. The tree was inferred using the 463 core gene families present in *Dickeya* and *Pectobacterium* strains. The scale bar represents the average number of substitutions; values at branch represent bootstrap values.

To further analyze the relatedness of *D. colocasiae* with other species of *Dickeya*, synteny analyses (Zhang et al., 2018; Pédron and Van Gijsegem, 2019) were performed with six available genomes. Genome synteny was visualized by a dot plot using the CLC Genomics Workbench v.20 (QIAGEN Bioinformatics, Aarhus, Denmark). The high synteny was conserved between the PL65^T^ and CE1 and showed collinearity with only a few inversions of genomic regions (**Figure 6A**). Although the inverted regions were larger, the syntenic regions were still dense between *D. zeae* Ech586 and PL65^T^ (**Figure 6B**). Comparison between *D. zeae* NCPPB 2538^T^ and PL65^T^ produced similar results (**Figure 6C**). In contrast, the syntenic regions between ZYY5^T^ *D. oryzae* and PL65^T^ were scattered (**Figure 6D**).

**Figure 6.**
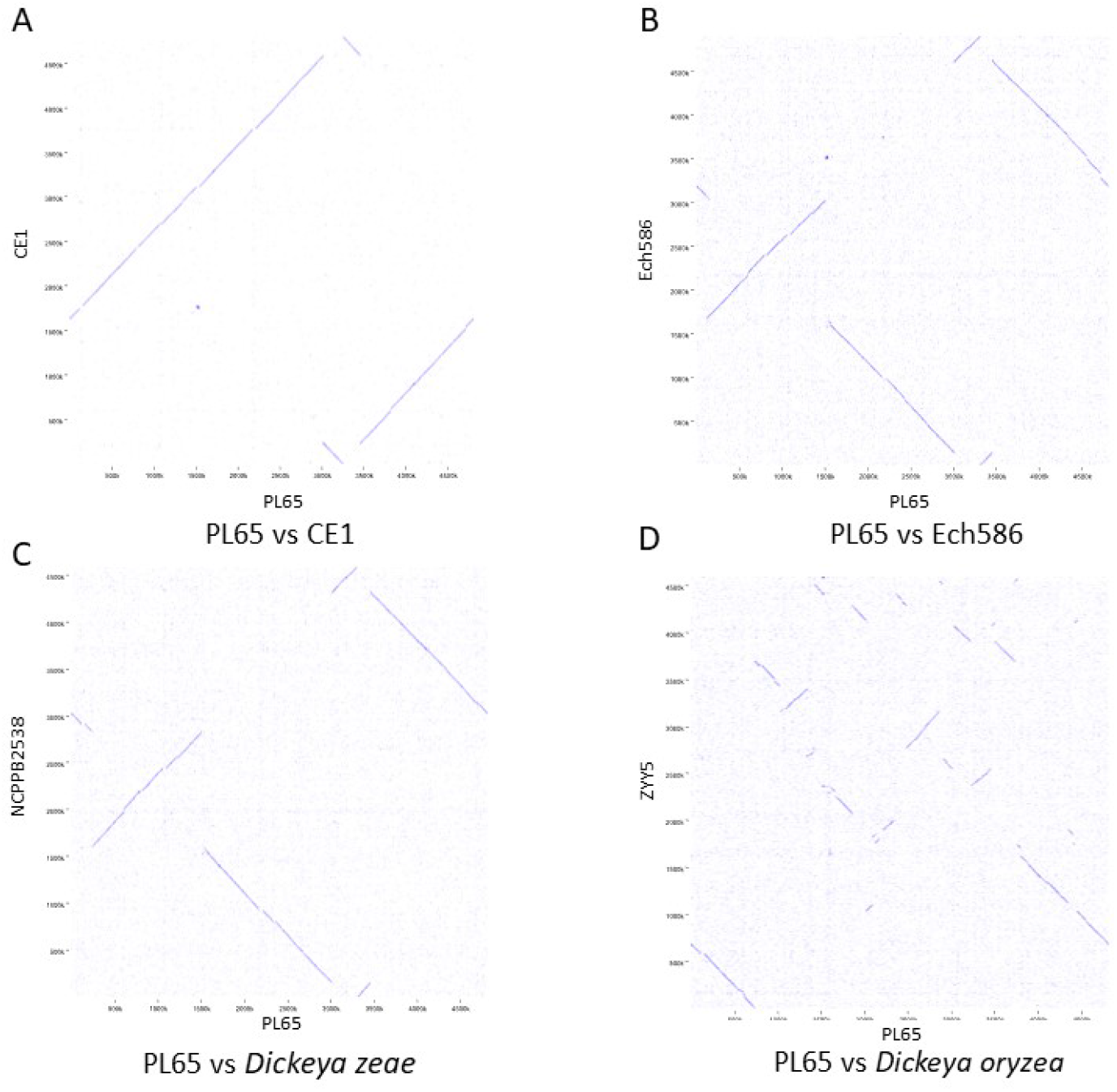
Synteny between *Dickeya colocasiae* and the other *Dickeya* strains from different species.

## Phenotypic Characteristics

Physiological and biochemical tests were performed for PL65^T^, A5432, and strains of eight closely related species: *Dickeya aquatica* (LGM 27354^T^), *D. dadantii* (CFBP 1269^T^ and CFBP 2051^T^), *D. dianthicola* (CFBP 1200^T^), *D. paradisiaca* (CFBP 4178^T^), *D. solani* (LMG 25993^T^), *D. zeae* (CFBP 2052^T^) and *D. chrysanthemi* (CFBP 2048^T^) using BIOLOG GEN III plates. The carbon source utilization was determined using Biolog GEN III MicroPlates according to the manufacturer’s instructions. Bacterial suspensions were prepared in inoculating fluid (IF) by adding pure bacterial colonies from the overnight grown culture on BUG (Biolog Universal Growth) media. The bacterial suspension was adjusted to achieve a 96% transmittance using a spectrophotometer SPECTRONIC 20D+ (Thermo Fisher Scientific, MA, USA). A 100 μL suspension was dispensed into each well of Biolog GEN III MicroPlate. The GEN III MicroPlates were incubated at 28 °C for 24h. The GEN III MicroPlates enable testing of Gram-negative and Gram-positive bacteria with 71 different carbon sources and 23 chemical sensitivity assays. The GEN III analyzes the ability of the cell to metabolize all major classes of the compounds, in addition to determining other important physiological properties, such as pH, salt and lactic acid tolerance, reducing power, and chemical sensitivity (**Table S3**). Although stachyose, inosine, and Tween 40 were utilized by the *D. zeae* type strain, the *D. colocasiae* strain showed no metabolic activities for these compounds. In contrast, the *Dickeya colocasiae* strains tolerated acetoacetic acid but *D. zeae* did not. (**Table 3**).

**Table 3.** Differential characteristics of strain PL65^T^ and closely related type strains Strain: Strain: LMG25993^T^ (D. *solanĩ*), CFBP4178^T^ (D. *paradisiaca*), CFBP2048^T^ (D. *chrysanthemi*), LGM27354 ^T^ (D. *aquatica*), CFBP1200^T^ (D. *dianthicola*), CFBP1269^T^ and CFBP2051^T^ (D. *dadantii*) and CFBP2052^T^ (D. zeae) All strains are positive for acid production from glycerol, d-arabinose, l-arabinose, ribose, d-xylose, galactose, rhamnose, inositol, arbutin, aesculin, salicoside, cellobiose and raffinose. Substrates utilized as carbon sources are α-d-glucose, d-mannose, d-fructose, glycerol, d-glucose-6-PO4, d-fructose-6-PO4, d-aspartic acid, l-aspartic acid, d-galacturonic acid, l-galactonic acid, lactone, mucic acid, d-saccharic acid, methyl pyruvate, citric acid, d-malic acid, l-malic acid, bromosuccinic acid and formic acid. All strains are negative for acid production from sorbitol, maltose, glycogen, d-lyxose, d-tagatose, d-fucose, lfucose, d-arabitol, l-arabitol and 5-keto-gluconate. Substrates that cannot be utilized as carbon sources are dextrin, maltose, trehalose, stachyose, N-acetyl neuraminic acid, d-serine, gelatin, glycyl-l-proline, p-hydroxy-phenylacetic acid, d-lactic acid methyl ester, γ-aminobutryric acid, α-hydroxy-butyric acid, β-hydroxy-d,l-butyric acid and α-keto-butyric acid. +/-, weakly utilized +, utilized; -, unutilized.

| Sugar Source Utilization | LMG25993<sup>T</sup> | CFBP4178<sup>T</sup> | CFBP2048<sup>T</sup> | LGM27354<sup>T</sup> | CFBP1200<sup>T</sup> | CFBP1269<sup>T</sup> | CFBP2051<sup>T</sup> | CFBP2052<sup>T</sup> | A5432 | PL65<sup>T</sup> |
| --- | --- | --- | --- | --- | --- | --- | --- | --- | --- | --- |
| Stachyose | - | +/- | +/- | - | - | - | - | +/- | - | - |
| Inosine | +/- | +/- | - | - | +/- | - | - | +/- | - | - |
| Dextrin | +/- | +/- | +/- | +/- | +/- | +/- | - | +/- | +/- | +/- |
| D-Maltose | - | - | - | +/- | - | +/- | - | +/- | +/- | +/- |
| D-Trehalose | +/- | - | +/- | +/- | - | +/- | - | +/- | +/- | +/- |
| D-Cellobiose | + | +/- | +/- | +/- | +/- | + | + | +/- | +/- | +/- |
| Gentiobiose | + | + | +/- | +/- | + | + | + | +/- | +/- | +/- |
| D-Turanose | +/- | - | +/- | +/- | - | - | - | +/- | +/- | +/- |
| α-D-Lactose | +/- | - | +/- | +/- | - | +/- | - | +/- | +/- | +/- |
| 3-methyl Glucose | - | - | - | - | - | - | - | +/- | +/- | +/- |
| D-Fucose | - | - | - | - | - | - | - | +/- | +/- | +/- |
| L-Fucose | - | - | +/- | - | - | - | - | +/- | +/- | +/- |
| L-Rhamnose | +/- | - | +/- | +/- | - | - | - | +/- | +/- | +/- |
| Sucrose | +/- | + | +/- | + | + | +/- | + | + | + | + |
| D-Raffinose | + | + | + | + | + | + | - | + | + | + |
| D-Mellobiose | + | + | +/- | + | + | + | - | +/- | + | + |
| D-Arabitol | +/- | - | - | - | - | - | - | +/- | +/- | +/- |
| 8% Nacl | - | - | - | - | - | - | +/- | - | - | - |
| 4% Nacl | +/- | - | +/- | +/- | - | +/- | + | +/- | +/- | +/- |
| D-Glucuronic acid | + | + | +/- | - | + | - | - | + | +/- | + |
| T-teen 40 | - | - | - | - | - | - | +/- | +/- | - | - |
| Acetoacetic acid | - | +/- | +/- | - | +/- | - | +/- | - | +/- | +/- |
| Fusidic acid | +/- | +/- | +/- | - | +/- | +/- | +/- | +/- | +/- | + |
| Lincomycin HCl | + | +/- | +/- | + | +/- | + | + | +/- | +/- | + |
| Gelatin | +/- | - | +/- | - | - | - | - | +/- | +/- | +/- |
| Glycyl-L-proline | +/- | - | +/- | - | - | + | - | +/- | +/- | +/- |
| L-Alanine | +/- | +/- | +/- | +/- | - | +/- | - | +/- | +/- | +/- |
| L-Arginine | - | - | +/- | - | - | - | - | +/- | +/- | +/- |
| L-Histidine | - | - | +/- | - | - | - | - | +/- | +/- | +/- |
| L-Pyroglutamic acid | - | - | +/- | - | - | - | - | +/- | +/- | +/- |
| L-Glutamic acid | +/- | +/- | +/- | + | +/- | + | - | + | +/- | + |
| pH 5 | +/- | +/- | - | - | - | - | +/- | - | - | - |
| pH 6 | + | + | +/- | + | + | + | + | + | + | + |
| Guanidine | + | - | +/- | +/- | - | +/- | + | +/- | +/- | +/- |

## DESCRIPTION OF *DICKEYA COLOCASIAE* SP. NOV

*Dickeya colocasiae* (co.lo.ca’si.ae L. fem. gen. n. colocasiae of taro)

The strain is gram-negative, motile and pectinolytic bacterium, producing pits on the crystal violet pectate (CVP) medium. Forms white, medium, and flat colonies on Nutrient Agar supplement with 0.4% glucose. Able to grow at 18, 28 and 32°C with an optimal growth temperature of 28°C on Nutrient Agar supplemented with 0.4% glucose. Colonies are smooth, irregular shaped with undulate margins, moist consistency, umbonate relief, bright yellow in color, when grown on solid SOB medium (2% tryptone, 0.5 % yeast extract, 10 mM NaCl, 2.5 mM KCl, 10 mM MgSO4, 1.5 % agar supplemented with 2 % glycerol) after 3 days at 28°C. The strain can cause soft rot on corms of taro plants under storage conditions and can survive in irrigation water. The strain is sensitive to bacitracin, carbenicillin, chloramphenicol, gentamicin, kanamycin, penicillin, tetracycline, minocycline, and aztreonam, and resistant to troleandomycin, rifamycin SV, fusidic acid, lithium chloride, sodium bromate, and potassium tellurite. It tolerates medium salinity up to 4 % NaCl and pH 6, and is resistant to 1 % sodium lactate, guanidine HCl, niaproof 4, tetrazolium violet, tetrazolium blue, and sodium butyrate. The strain was able to consume comparatively more sugars and amino acids; sucrose, D-raffinose, D-mellobiose, B-methyl-D-glucoside, D-salicin, N-acetyl-D-glucosamine, α-D-glucose, D-mannose, D-fructose, D-galactose, D-sorbitol, D-mannitol, myo-inositol, glycerol, D-glucose-6-PO4, D-fructose-6-PO4, D-aspartic acid, L-aspartic acid, L-glutamic acid, L-serine, pectin, D-galacturonic acid, L-galactonic acid, lactone, D-gluconic acid, D-glucuronic acid, mucic acid, D-saccharic acid, methyl pyruvate, L-lactic acid, citric acid, D-malic acid, L-malic acid, bromo-succinic acid, acetic acid, and formic acid. Able to weakly utilize dextrin, D-maltose, D-trehalose, D-cellobiose, gentiobiose, D-turanose α-D-lactose, N-acetyl-β-D-mannosamine, N-acetyl-D-galactosamine, 3-methyl glucose, D-fucose, L-fucose, L-rhamnose, D-arabitol, gelatin, glycyl-L-proline, L-alanine, L-arginine, L-histidine, L-pyroglutamic acid, glucuronamide, and acetoacetic acid. Unable to utilize stachyose, N-acetyl-neuraminic acid, inosine, D-serine, quinic acid, p-hydroxy-phenylacetic acid, D-lactic acid, methyl ester, A-keto-glutaric acid, Tween 40, γ-aminobutyric acid, α-hydroxy-butyric acid, β-hydroxy-D, L-butyric acid, α-keto-butyric acid, and propionic acid.

Bacteria in this species have been isolated from taro plants expressing soft rot symptoms after harvest and from irrigation water in Hawaii. The type strain is PL65^T^ (ICMP 24361^T^). The DNA G+C content of the type strain is 53.6 based on the completed genome sequence.

## AUTHOR STATEMENTS

### Ethical approval

*N/A*

### Consent for publication

*N/A*

