## Supplementary material for "*Dickeya colocasiae* sp. nov. isolated from wetland taro, *Colocasia esculentum*": Supplemet Data

### SUPPLEMENTARY DATA

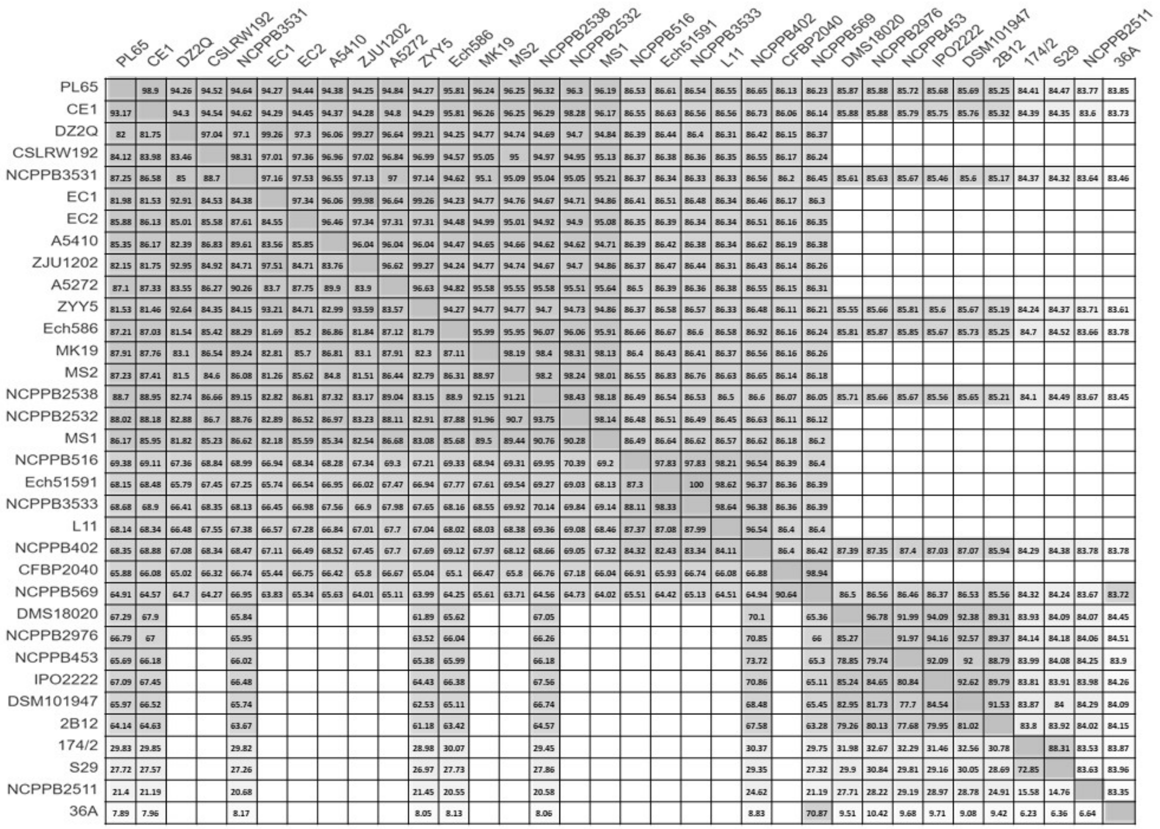


**Figure S1.** Pairwise heatmap based on the average nucleotide identity (ANIm) and the average nucleotide coverage (or Alignment percentage; AP). The ANIm (upper triangle) and AP (lower triangle) values for strain PL65 and other *Dickeya* strains with available genomes. Darker colors used to represent higher values.

**Table S1**. The sequences of all *Dickeya* strains, *Pectobacterium atrosepticum* and *Pectobacterium brasiliense* used for multi-locus sequence analysis (MLSA) study. Type strains are marked with “$^{T}$” after the strain name. The data not available is marked as “-”.

| Species | Strain | Accession Number | | | | |
|---|---|---|---|---|---|---|
| | | ***dnaA*** | ***gapA*** | ***gyrB*** | ***atpD*** | ***purA*** |
| *Dickeya colocasiae* | CE1 | NZ_CP033622.1 | NZ_CP033622.1 | NZ_CP033622.1 | NZ_CP033622.1 | NZ_CP033622.1 |
| *Dickeya colocasiae* | PL65$^{T}$ | NZ_CP040817.1 | NZ_CP040817.1 | NZ_CP040817.1 | NZ_CP040817.1 | NZ_CP040817.1 |
| *Dickeya colocasiae* | A5432 | MW791166 | MW791117 | MT017771 | MW790982 | MW791050 |
| *Dickeya colocasiae* | A5511 | MW791170 | MW791113 | MT017767 | MW790978 | MW791046 |
| *Dickeya colocasiae* | A5612 | MW791173 | MW791106 | MT017760 | MW790971 | MW791039 |
| *Dickeya colocasiae* | A5428 | MW791165 | MW791118 | MT017772 | MW790983 | MW791051 |
| *Dickeya aquatica* | 174/2$^{T}$ | NZ_LT615367.1 | NZ_LT615367.1 | NZ_LT615367.1 | NZ_LT615367.1 | NZ_LT615367.1 |
| *Dickeya chrysanthemi* | Ech1591 | NC_012912.1 | NC_012912.1 | NC_012912.1 | NC_012912.1 | NC_012912.1 |
| *Dickeya chrysanthemi* | L11 | NZ_JSYH01000045.1 | NZ_JSYH01000005.1 | NZ_JSYH01000045.1 | NZ_JSYH01000064.1 | NZ_JSYH01000039.1 |
| *Dickeya chrysanthemi* | NCPPB 3533 | NZ_CM001981.1 | NZ_CM001981.1 | NZ_CM001981.1 | NZ_CM001981.1 | NZ_CM001981.1 |
| *Dickeya chrysanthemi* | NCPPB 402$^{T}$ | NZ_CM001974.1 | NZ_CM001974.1 | NZ_CM001974.1 | NZ_CM001974.1 | NZ_CM001974.1 |
| *Dickeya chrysanthemi* | NCPPB 516 | NZ_CM001904.1 | NZ_CM001904.1 | NZ_CM001904.1 | NZ_CM001904.1 | NZ_CM001904.1 |
| *Dickeya dadantii* | DSM 18020$^{T}$ | NZ_CP023467.1 | NZ_CP023467.1 | NZ_CP023467.1 | NZ_CP023467.1 | NZ_CP023467.1 |
| *Dickeya dadantii* | NCPPB 2976$^{T}$ | NZ_CM001978.1 | NZ_CM001978.1 | NZ_CM001978.1 | NZ_CM001978.1 | NZ_CM001978.1 |
| *Dickeya dianthicola* | NCPPB 453$^{T}$ | NZ_CM001841.1 | NZ_CM001841.1 | NZ_CM001841.1 | NZ_CM001841.1 | NZ_CM001841.1 |
| *Dickeya fangzhongdai* | DSM 101947$^{T}$ | NZ_CP025003.1 | NZ_CP025003.1 | NZ_CP025003.1 | NZ_CP025003.1 | NZ_CP025003.1 |
| *Dickeya lacustris* | S29$^{T}$ | NZ_QNUT01000058.1 | NZ_QNUT01000032.1 | NZ_QNUT01000058.1 | NZ_QNUT01000058.1 | NZ_QNUT01000054.1 |
| *Dickeya oryzae* | CSL RW192 | NZ_CM001972.1 | NZ_CM001972.1 | NZ_CM001972.1 | NZ_CM001972.1 | NZ_CM001972.1 |
| *Dickeya oryzae* | DZ2Q | NZ_APWM01000004.1 | NZ_APWM01000009.1 | NZ_APWM01000004.1 | NZ_APWM01000004.1 | NZ_APWM01000001.1 |
| *Dickeya oryzae* | EC1 | NZ_CP006929.1 | NZ_CP006929.1 | NZ_CP006929.1 | NZ_CP006929.1 | NZ_CP006929.1 |
| *Dickeya oryzae* | EC2 | NZ_CP031515.1 | NZ_CP031515.1 | NZ_CP031515.1 | NZ_CP031515.1 | NZ_CP031515.1 |
| *Dickeya oryzae* | NCPPB 3531 | NZ_CM001980.1 | NZ_CM001980.1 | NZ_CM001980.1 | NZ_CM001980.1 | NZ_CM001980.1 |
| *Dickeya oryzae* | ZJU1202 | NZ_AJVN01000004.1 | NZ_AJVN01000003.1 | NZ_AJVN01000004.1 | NZ_AJVN01000004.1 | NZ_AJVN01000001.1 |

| | | | | | | |
|---|---|---|---|---|---|---|
| *Dickeya oryzae* | ZYY5 $^{T}$ | NZ_SZVX01000007.1 | NZ_SZVX01000001.1 | NZ_SZVX01000007.1 | NZ_SZVX01000007.1 | NZ_SZVX01000016.1 |
| *Dickeya oryzea* | A5272 | NZ_CP040816.1 | NZ_CP040816.1 | NZ_CP040816.1 | NZ_CP040816.1 | NZ_CP040816.1 |
| *Dickeya oryzea* | A5410 | NZ_CP040817.1 | NZ_CP040817.1 | NZ_CP040817.1 | NZ_CP040817.1 | NZ_CP040817.1 |
| *Dickeya paradisiaca* | NCPPB 2511 $^{T}$ | NZ_CM001857.1 | NZ_CM001857.1 | NZ_CM001857.1 | NZ_CM001857.1 | NZ_CM001857.1 |
| *Dickeya poaceiphila* | NCPPB 569 $^{T}$ | NZ_CP042220.2 | NZ_CP042220.2 | NZ_CP042220.2 | NZ_CP042220.2 | NZ_CP042220.2 |
| *Dickeya poaceiphila* | CFBP 2040 | NZ_JAAVXH010000074.1 | NZ_JAAVXH010000120.1 | NZ_JAAVXH010000074.1 | NZ_JAAVXH010000074.1 | NZ_JAAVXH010000087.1 |
| *Dickeya solani* | IPO 2222 $^{T}$ | NZ_CP015137.1 | NZ_CP015137.1 | NZ_CP015137.1 | NZ_CP015137.1 | NZ_CP015137.1 |
| *Dickeya undicola* | 2B12 $^{T}$ | NZ_JSYG01000008.1 | NZ_JSYG01000031.1 | NZ_JSYG01000008.1 | NZ_JSYG01000008.1 | NZ_JSYG01000001.1 |
| *Dickeya zeae* | Ech586 | NC_013592.1 | NC_013592.1 | NC_013592.1 | NC_013592.1 | NC_013592.1 |
| *Dickeya zeae* | MK19 | NZ_CM001985.1 | NZ_CM001985.1 | NZ_CM001985.1 | NZ_CM001985.1 | NZ_CM001985.1 |
| *Dickeya zeae* | MS1 | NZ_APMV01000026.1 | NZ_APMV01000015.1 | NZ_APMV01000026.1 | NZ_APMV01000025.1 | NZ_APMV01000024.1 |
| *Dickeya zeae* | MS2 | NZ_CP025799.1 | NZ_CP025799.1 | NZ_CP025799.1 | NZ_CP025799.1 | NZ_CP025799.1 |
| *Dickeya zeae* | NCPPB 2538 $^{T}$ | NZ_CM001977.1 | NZ_CM001977.1 | NZ_CM001977.1 | NZ_CM001977.1 | NZ_CM001977.1 |
| *Dickeya zeae* | NCPPB 3532 | NZ_CM001858.1 | NZ_CM001858.1 | NZ_CM001858.1 | NZ_CM001858.1 | NZ_CM001858.1 |
| *Pectobacterium atrosepticum* | 36A | NZ_CP024956.1 | NZ_CP024956.1 | NZ_CP024956.1 | NZ_CP024956.1 | NZ_CP024956.1 |
| *Pectobacterium brasiliense* | BC1 | NZ_CP009769.1 | NZ_CP009769.1 | NZ_CP009769.1 | NZ_CP009769.1 | NZ_CP009769.1 |

**Table S2**. The information about the *Dickeya* species and *Pectobacterium atrosepticum* genome sequences used for core genes study. Type strains are marked with “[T]” after the strain name. The data not available is marked with “-”.

| Species | Strain | Accession Number | Location | Host/Source | Isolation Year | Size (Mb) | GC % |
|---|---|---|---|---|---|---|---|
| *Dickeya new* | CE1 | NZ_CP033622 | China | *Canna edulis* | 2017 | 4.71473 | 53.6 |
| *Dickeya new* | PL65 [T] | NZ_CP040817 | USA | *Colocasia esculenta* | 2018 | 4.749968 | 53.6 |
| *Dickeya aquatica* | 174/2 [T] | NZ_LT615367 | UK | River water | 2012 | 4.50156 | 53.6 |
| *Dickeya chrysanthemi* | Ech1591 | NC_012912 | - | *Zea mays* | - | 4.81385 | 54.5 |
| *Dickeya chrysanthemi* | L11 | NZ_JSYH0100 | Malaysia | Lake water | 2014 | 4.76788 | 54.3 |
| *Dickeya chrysanthemi* | NCPPB 3533 | NZ_CM001981 | USA | *Solanum tuberosum* | 1985 | 4.73071 | 54.51 |
| *Dickeya chrysanthemi* | NCPPB 402 [T] | NZ_CM001974 | USA | *Chrysanthemum morifolium* | 1956 | 4.70035 | 54.19 |
| *Dickeya chrysanthemi* | NCPPB 516 | NZ_CM001904 | Denmark | *Parthenium argentatum* | 1957 | 4.61724 | 54.24 |
| *Dickeya dadantii* | DSM 18020 [T] | NZ_CP023467 | Comoro Islands | *Pelargonium capitatum* | 1960 | 4.99754 | 56.4 |
| *Dickeya dadantii* | NCPPB 2976 [T] | NZ_CM001978 | USA | *Dieffenbachia sp.* | 1957 | 4.81753 | 56.41 |
| *Dickeya dadantii* | 3937 | NC_014500 | France | *Saintpaulia ionantha* | 1977 | 4.9228 | 56.3 |
| *Dickeya dianthicola* | NCPPB 453 [T] | NZ_CM001841 | UK | *Dianthus caryophyllus* | 1956 | 4.67606 | 55.95 |
| *Dickeya dianthicola* | ME23 | NZ_CP031560 | USA Maine | *S. tuberosum* | 2016 | 4.90906 | 55.7 |
| *Dickeya fangzhongdai* | DSM 101947 [T] | NZ_CP025003 | China | *Pyrus pyrifolia* | 2009 | 5.03245 | 56.79 |
| *Dickeya fangzhongdai* | PA1 | NZ_CP020872 | China | *Phalaenopsis* sp. | 2011 | 4.97922 | 56.9 |
| *Dickeya lacustris* | S29 [T] | NZ_QNUT01 | France | *River water* | 2017 | 4.306505 | 53.1 |
| *Dickeya oryzae* | CSL RW192 | NZ_CM001972 | UK | *River water* | 2012 | 4.70177 | 53.39 |
| *Dickeya oryzae* | DZ2Q | NZ_APWM01 | Italy | *Oryza sativa* | - | 4.64652 | 53.2 |
| *Dickeya oryzae* | EC1 | NZ_CP006929 | China | *Oryza sativa* | - | 4.53236 | 53.4 |
| *Dickeya oryzae* | EC2 | NZ_CP031515 | China | *Oryza sativa* | 2016 | 4.57512 | 53.3 |
| *Dickeya oryzae* | NCPPB 3531 | NZ_CM001980 | Australia | *S. tuberosum* | 1987 | 4.62586 | 53.7 |
| *Dickeya oryzae* | ZJU1202 | NZ_AJVN0100 | China | *Oryza sativa* | 2012 | 4.58711 | 53.3 |
| *Dickeya oryzae* | ZYY5 [T] | NZ_SULL00 | China | *Oryza sativa* | 2010 | 4.58718 | 53.3 |
| *Dickeya oryzea* | A5272 | NZ_CP040816 | USA | *Ananas comosus* | 2003 | 4.697973 | 53.6 |
| *Dickeya oryzea* | A5410 | NZ_CP040817 | USA | *Ananas comosus* | 2007 | 4.779199 | 53.5 |
| *Dickeya paradisiaca* | NCPPB 2511 [T] | NZ_CM001857 | Colombia | *M. paradisiaca* var. *dominico* | 1970 | 4.63187 | 55 |
| *Dickeya paradisiaca* | Ech703 | NC_012880 | Australia | *S. tuberosum* | - | 4.67945 | 55 |

| | | | | | | | |
|---|---|---|---|---|---|---|---|
| *Dickeya poaceiphila* | NCPPB 569 $^{T}$ | NZ_CP042220 | Australia | *Saccharum officinarum* | 1958 | 4.31715 | 52.7 |
| *Dickeya poaceiphila* | CFBP 2040 | NZ_JAAVXH01 | Australia | *Megathyrsus maximus* | 1980 | 4.0214 | 52.8 |
| *Dickeya solani* | IPO 2222 $^{T}$ | NZ_CP015137 | Netherland | *S. tuberosum* | 2007 | 4.91983 | 56.2 |
| *Dickeya undicola* | 2B12 $^{T}$ | NZ_JSYG01 | Malaysia | Fresh water | 2014 | 4.34979 | 54.5 |
| *Dickeya undicola* | FVG10-MFV-A16 | NZ_RJLS00 | France | Fresh water | 2017 | 4.54214 | 54.5 |
| *Dickeya zeae* | Ech586 | NC_013592 | USA Florida | *Philodendron Schott* | - | 4.81839 | 53.6 |
| *Dickeya zeae* | MK19 | NZ_CM001985 | Scotland | River water | 2012 | 4.67215 | 53.6 |
| *Dickeya zeae* | MS1 | NZ_APMV0100 | China | *Musa* sp. | 2009 | 4.74828 | 53.3 |
| *Dickeya zeae* | MS2 | NZ_CP025799 | China | *Musa* sp. | 2014 | 4.74005 | 53.4 |
| *Dickeya zeae* | NCPPB 2538 $^{T}$ | NZ_CM001977 | USA | *Zea mays* | 1970 | 4.56379 | 53.6 |
| *Dickeya zeae* | NCPPB 3532 | NZ_CM001858 | Australia | *S. tuberosum* | 1987 | 4.55706 | 53.6 |
| *Pectobacterium atrosepticum* | 36A | NZ_CP024956 | Belarus | *S. tuberosum* | 1978 | 4.96558 | 51.1 |

**Table S3**. Phenotypic characters of *Dickeya colocasiae* strain PL65$^{T}$ and A5432 and closely related type strains of *Dickeya* species by using GEN III MicroPlates. Strains: LMG25993$^{T}$ (*D. solani*), CFBP4178$^{T}$ (*D. paradisiaca*), CFBP2048$^{T}$ (*D. chrysanthemi*), LGM27354 $^{T}$ (*D. aquatica*), CFBP1200$^{T}$ (*D. dianthicola*), CFBP1269$^{T}$ and CFBP2051$^{T}$ (*D. dadantii*) and CFBP2052$^{T}$ (*D. zeae*)

| Sugar Source Utilization | LMG25993$^{T}$ | CFBP 4178$^{T}$ | CFBP 2048$^{T}$ | LGM27354$^{T}$ | CFBP 1200$^{T}$ | CFBP1269$^{T}$ | CFBP2051$^{T}$ | CFBP2052$^{T}$ | A5432 | PL65$^{T}$ |
|---|---|---|---|---|---|---|---|---|---|---|
| **Dextrin** | +/- | +/- | +/- | +/- | +/- | +/- | - | +/- | +/- | +/- |
| **D-Maltose** | - | - | - | +/- | - | +/- | - | +/- | +/- | +/- |
| **D-Trehalose** | +/- | - | +/- | +/- | - | +/- | - | +/- | +/- | +/- |
| **D-Cellobiose** | + | +/- | +/- | +/- | +/- | + | + | +/- | +/- | +/- |
| **Gentiobiose** | + | + | +/- | +/- | + | + | + | +/- | +/- | +/- |
| **Sucrose** | +/- | + | +/- | + | + | +/- | + | + | + | + |
| **D-Turanose** | +/- | - | +/- | +/- | - | - | - | +/- | +/- | +/- |
| **Stachyose** | - | +/- | +/- | - | - | - | - | +/- | - | - |
| **pH 6** | + | + | +/- | + | + | + | + | + | + | + |
| **pH 5** | +/- | +/- | - | - | - | - | +/- | - | - | - |
| **D-Raffinose** | + | + | + | + | + | + | - | + | + | + |
| **α-D-Lactose** | +/- | - | +/- | +/- | - | +/- | - | +/- | +/- | +/- |
| **D-Mellobiose** | + | + | +/- | + | + | + | - | +/- | + | + |
| **B-Methyl-D-Glucoside** | + | + | + | + | + | + | + | + | + | + |
| **D-Salicin** | + | + | +/- | + | + | + | + | + | + | + |
| **N-acetyl-D-Glucosamine** | + | + | + | + | + | + | + | + | + | + |

| | | | | | | | | | | |
|---|---|---|---|---|---|---|---|---|---|---|
| **N-acetyl-β-D-Mannosamine** | +/- | +/- | +/- | +/- | +/- | +/- | - | +/- | +/- | +/- |
| **N-acetyl-D-Galactosamine** | - | - | - | +/- | - | - | - | +/- | +/- | +/- |
| **N-acetyl-Neuraminic acid** | - | - | - | - | - | - | - | - | - | - |
| **1% Nacl** | + | + | + | + | + | + | + | + | + | + |
| **4% Nacl** | +/- | - | +/- | +/- | - | +/- | + | +/- | +/- | +/- |
| **8% Nacl** | - | - | - | - | - | - | +/- | - | - | - |
| **α-D-Glucose** | + | + | +/- | + | + | + | + | + | + | + |
| **D-Mannose** | + | + | + | + | + | + | + | + | + | + |
| **D-Fructose** | + | + | + | + | + | + | + | + | + | + |
| **D-Galactose** | + | + | + | + | + | + | + | + | + | + |
| **3-methyl Glucose** | - | - | - | - | - | - | - | +/- | +/- | +/- |
| **D-Fucose** | - | - | - | - | - | - | - | +/- | +/- | +/- |
| **L-Fucose** | - | - | +/- | - | - | - | - | +/- | +/- | +/- |
| **L-Rhamnose** | +/- | - | +/- | +/- | - | - | - | +/- | +/- | +/- |
| **Inosine** | +/- | +/- | - | - | +/- | - | - | +/- | - | - |
| **1% Sodium lactate** | + | + | + | + | + | + | + | + | + | + |
| **Fusidic acid** | +/- | +/- | +/- | - | +/- | +/- | +/- | +/- | +/- | + |
| **D -Serine** | - | - | - | - | +/- | - | +/- | - | - | - |
| **D-Sorbitol** | +/- | - | +/- | - | - | + | - | +/- | + | + |
| **D-Mannitol** | + | - | +/- | - | - | +/- | + | + | + | + |

| | | | | | | | | | | |
|---|---|---|---|---|---|---|---|---|---|---|
| **D-Arabitol** | +/- | - | - | - | - | - | - | +/- | +/- | +/- |
| **myo-Inositol** | + | - | +/- | + | - | + | + | + | + | + |
| **Glycerol** | + | + | + | + | + | + | + | + | + | + |
| **D-Glucose-6-PO4** | + | + | + | + | + | + | + | + | + | + |
| **D-Fructose-6-PO4** | + | + | + | + | + | + | + | + | + | + |
| **D-Aspartic acid** | + | +/- | +/- | + | + | + | + | + | + | + |
| **D-Serine** | - | - | - | - | - | - | - | - | - | - |
| **Troleandomycin** | + | + | + | + | + | + | + | + | + | + |
| **Rifamycin SV** | + | + | + | + | + | + | + | + | + | + |
| **Minocycline** | - | - | - | - | - | - | - | - | - | - |
| **Gelatin** | +/- | - | +/- | - | - | - | - | +/- | +/- | +/- |
| **Glycyl-L-proline** | +/- | - | +/- | - | - | + | - | +/- | +/- | +/- |
| **L-Alanine** | +/- | +/- | +/- | +/- | - | +/- | - | +/- | +/- | +/- |
| **L-Arginine** | - | - | +/- | - | - | - | - | +/- | +/- | +/- |
| **L-Aspartic acid** | + | + | + | + | + | + | + | + | + | + |
| **L-Glutamic acid** | +/- | +/- | +/- | + | +/- | + | - | + | +/- | + |
| **L-Histidine** | - | - | +/- | - | - | - | - | +/- | +/- | +/- |
| **L-Pyroglutamic acid** | - | - | +/- | - | - | - | - | +/- | +/- | +/- |
| **L-Serine** | + | + | + | + | + | + | - | + | + | + |
| **Lincomycin HCl** | + | +/- | +/- | + | +/- | + | + | +/- | +/- | + |

| | | | | | | | | | | |
|---|---|---|---|---|---|---|---|---|---|---|
| **Guanidine** | + | - | +/- | +/- | - | +/- | + | +/- | +/- | +/- |
| **Niaproof 4** | + | + | + | + | + | + | + | + | + | + |
| **Pectin** | + | + | + | + | + | + | + | + | + | + |
| **D-Galacturonic acid** | + | + | + | + | + | + | + | + | + | + |
| **L-Galactonic acid lactone** | + | + | + | + | + | + | + | + | + | + |
| **D-Gluconic acid** | + | + | +/- | + | + | + | + | + | + | + |
| **D-Glucuronic acid** | + | + | +/- | - | + | - | - | + | +/- | + |
| **Glucuronamide** | +/- | +/- | +/- | +/- | +/- | +/- | +/- | +/- | +/- | +/- |
| **Mucic acid** | + | + | + | + | + | + | + | + | + | + |
| **Quinic acid** | - | - | - | - | - | - | - | - | - | - |
| **D-Saccharic acid** | + | + | + | + | + | + | + | + | + | + |
| **Vancomycin** | + | + | + | + | + | + | + | + | + | + |
| **Tetrazolium violet** | + | +/- | + | + | +/- | + | + | + | + | + |
| **Tetrazolium blue** | + | +/- | + | + | + | + | + | + | + | + |
| **p-Hydroxy-phenylacetic acid** | - | - | - | - | - | - | - | - | - | - |
| **Methyl pyruvate** | + | + | + | + | + | + | + | + | + | + |
| **D-Lactic acid methyl ester** | - | - | - | - | - | - | - | - | - | - |
| **L-Lactic acid** | + | + | +/- | - | + | + | + | + | + | + |
| **Citric acid** | + | + | + | + | + | + | + | + | + | + |
| **A-keto-Glutaric acid** | - | - | - | - | - | - | - | - | - | - |

| | | | | | | | | | | |
|---|---|---|---|---|---|---|---|---|---|---|
| **D-Malic acid** | + | + | - | + | + | + | + | + | + | + |
| **L-Malic acid** | + | + | + | + | + | + | + | + | + | + |
| **Bromo-succinic acid** | + | + | +/- | + | +/- | + | + | + | + | + |
| **Nalidixic acid** | - | - | - | - | - | - | - | - | - | - |
| **Lithium chloride** | + | - | + | +/- | - | + | + | + | + | + |
| **Potassium tellurite** | - | - | - | + | - | +/- | - | - | - | - |
| **Tween 40** | - | - | - | - | - | - | +/- | +/- | - | - |
| **γ-amino-Butyric acid** | - | - | - | - | - | - | - | - | - | - |
| **α-Hydroxy-butyric acid** | - | +/- | - | - | +/- | - | - | - | - | - |
| **β-Hydroxy-D, L-butyric acid** | - | - | - | - | - | - | - | - | - | - |
| **α-keto-Butyric acid** | - | - | - | - | - | - | - | - | - | - |
| **Acetoacetic acid** | - | +/- | +/- | - | +/- | - | +/- | - | +/- | +/- |
| **Propionic acid** | - | - | - | - | - | - | - | - | - | - |
| **Acetic acid** | +/- | +/- | +/- | + | + | + | + | + | + | + |
| **Formic acid** | + | +/- | +/- | +/- | +/- | +/- | + | + | + | + |
| **Aztreonam** | +/- | +/- | - | - | - | - | - | - | - | - |
| **Sodium butyrate** | +/- | + | - | +/- | +/- | +/- | +/- | +/- | +/- | +/- |
| **Sodium bromate** | - | - | - | - | - | - | - | - | - | - |
